# The effects of sex on extinction dynamics and evolutionary rescue of *Chlamydomonas reinhardtii* depend on the rate of environmental change

**DOI:** 10.1101/817056

**Authors:** Nikola Petkovic, Nick Colegrave

**Author notes:** **Corresponding author:** Nikola Petkovic, Koç University, Rumeli Feneri Yolu, Sarıyer, 33450, İstanbul, Turkey.

## Abstract

The continued existence of sex despite many costs it entails still lacks an adequate explanation. Previous experimental studies demonstrated that sex enhances the rate of adaptation in changing environments. To the best of our knowledge, no experimental study that investigated the effects of sex on adaptation has manipulated the rate of environmental change, which is negatively correlated with the probability of population survival. Since the patterns of adaptation (adaptive walk) depend on the rate of environmental change, the overall effects of sex may also be altered by this factor.

To investigate the interplay of sex and the rate of environmental deterioration, we carried out a long-term selection experiment with a unicellular alga (*Chlamydomonas reinhardtii)*, by manipulating mode of reproduction (asexual or sexual populations) and the rate of environmental deterioration (an increase of salt concentration). We monitored both the population size and extinction dynamics and estimated the probability of evolutionary rescue.

We detected a faster decline of population size in the asexual group relative to both the obligate sexual and facultative sexual group, irrespective of the rate of environmental deterioration. The results revealed significant interaction between mode of reproduction and the rate of environmental deterioration on extinction rate of experimental populations. Obligate sexual reproduction was advantageous under the intermediate rate of environmental deterioration, while facultative sexuality was favoured under the high rate of environmental deterioration. The populations within the obligate sexual group were most likely to adapt to grow in conditions lethal for the ancestral populations, irrespective of the rate of environmental deterioration.

To the best of our knowledge, this is the first study which indicates that different modes of sexual reproduction will be beneficial (slower extinction rate) at different rates of environmental deterioration and that obligate sexuality maximizes the probability of adaptation irrespective of the rate of environmental deterioration.

## 1. Introduction

Despite the fact that the great majority of extant eukaryotes are sexual, sex still remains one of the most intriguing puzzles in evolutionary biology (Otto and Lenormand, 2002). The reason for this reflects in the numerous disadvantages and costs of sex (Lewis, 1987; Lehtonen et al., 2012). These costs can be significant and hence the continued existence of sex implies large potential benefits. Unexpected difficulty to provide an explanation for such a widespread phenomenon brought sex the title of “the Queen of problems in evolutionary biology” (Bell, 1982).

Most contemporary hypotheses explaining the maintenance of sex by natural selection are based on the idea that sex increases genetic variability of the progeny (Kondrashov, 1993). This effect, in turn, increases the response of selection and thus, facilitates the rate of adaptation to novel environments (Weismann, 1889; Fisher, 1930; Muller, 1932; Burt, 2000). The advantageous effect of sex based on this concept is manifested through faster accumulation of mutations beneficial in changing conditions in sexual populations, which increases the rate of adaptation relative to asexual populations. In asexual populations, which may fix novel mutations at a slower rate (one at the time), adaptation proceeds at a lower rate or can be limited by competition of individuals carrying different mutations (termed clonal interference) (Muller 1932; Gerrish and Lenski, 1998).

A substantial body of experimental evidence indicates that sex increases the rate of adaptation to changing environments (Hartfield and Keightley, 2012). The advantageous effect of sex is manifested through faster assembling of mutations beneficial in novel environments (Kaltz and Bell, 2002), reduction of clonal interference in large populations (Colegrave, 2002), an increase in fitness of the progeny (Becks and Agrawal, 2012) and more efficient adaptation to the presence of pathogenic species (Morran et al., 2009). Furthermore, sex increases the rate of adaptation in a deteriorating environment (Goddard et al., 2005) and the probability of survival in conditions lethal for the ancestral population (Lachapelle and Bell, 2012), termed evolutionary rescue (Gomulkiewicz and Holt, 1995).

While the benefits of sex in novel and deteriorating environments are relatively well understood, to the best of our knowledge, no experimental studies which investigated the effects of sex have manipulated the rate of environmental change, which is negatively correlated with the probability of population survival in stressful environments (Perron et al., 2007; Bell and Gonzales, 2011; Ramsayer et al., 2013; Lindsey et al., 2013). Adaptation occurs through sequential fixation of beneficial mutations (adaptive walk) (Collins et al., 2007), a process which patterns depend on the rate of environmental change (Collins and de Meaux, 2009). Since sex directly impacts the adaptive walk through accumulation of beneficial mutations, the overall effects of it may be altered by different rates of environmental change.

The higher rates of environmental deterioration imply a larger initial maladaptedness of a population and adaptation may occur through strong selection, which causes a demographic pressure manifested through higher proportion of selective deaths (Haldane, 1957; Burger and Lynch, 1995). Furthermore, adaptation to the higher rates of environmental change occurs through fixation of few mutations of larger fitness effects (Collins and de Meaux, 2009), which are less frequent in the distribution of beneficial mutations (Kassen and Bataillon, 2006; Eyre-Walker and Keightley, 2007). Consequently, if population size substantially declines, the mutations which increase the fitness under the higher rates of environmental deterioration may appear in disadvantageous genetic background (with deleterious alleles) due to genetic drift (Otto and Lenormand, 2002). As a result, adaptation of an asexual population may become impeded and the population may face extinction. The potential beneficial effect of sex under these conditions is the release of these mutations from disadvantageous genetic background, which may restore the adaptive walk of the population. However, since demographic cost of adaptation imposes the limit for the rate of environmental change that a population can sustain (Lynch and Lande, 1993), the beneficial effect of sex may diminish if the magnitude of environmental change outweighs the reproductive capacity of a species to compensate for the demographic loss.

Adaptation to a gradual rate of environmental change occurs through fixation of mutations of smaller fitness effects, which are more frequent in the distribution of beneficial mutations (Eyre-Walker and Keightley, 2007). In addition, a population subjected to the gradual rate of environmental deterioration will likely suffer a less pronounced population size decline than a population subjected to the higher rate of environmental change and hence the supply of beneficial mutations will be proportionally higher. However, due to weaker selection operating under the gradual environmental change, these mutations may not reach fixation, or otherwise fix in less predictable intervals (a process termed periodic fixation) (Collins and de Meaux, 2009). If two or more mutations, which are beneficial under the gradual rates of environmental deterioration, segregate in an asexual population, adaptation may slow down due to clonal interference (Muller, 1932). The potential advantage of sex under the gradual rate of environmental deterioration may manifest through alleviation of clonal interference and thereby a higher rate of adaptation of sexual populations, as demonstrated during adaptation to novel environments (Colegrave, 2002).

In order to investigate the interplay of sex and the rate of environmental deterioration on adaptation to changing conditions, we carried out a selection experiment by subjecting the experimental populations of a unicellular alga *Chlamydomonas reinhardtii* to the environment which progressively deteriorated and ultimately became lethal for the ancestral populations. We manipulated both their mode of reproduction and the rate of environmental deterioration in a fully factorial design and monitored the extinction dynamics and the occurrence of evolutionary rescue events within the treatment groups.

We predicted that the sign of the effect of sex (slower extinction rate and a higher probability of evolutionary rescue of the sexual populations) will not depend on the rate of environmental deterioration, despite the potential differences in the patterns of the effects of sex under different rates of environmental change. Furthermore, we predicted that the asexual populations will adapt less efficiently under the higher rates of environmental change, hence the magnitude of the effect of sex (the relative advantage of sexual populations) will be positively correlated with the rate of environmental deterioration.

## 2. Materials and methods

### 2.1 Base populations of *C. reinhardtii*

To establish genetically variable experimental populations of *C. reinhardtii*, the mass mating of 10 different wild type isolates was carried out. The zygotes obtained were transferred to the agar plates by sterile loops and allowed to germinate by producing zoospores. The zoospores differentiated into adult vegetative cells, which were allowed to undergo several rounds of mitotic divisions (for 3-4 days) until visible colonies appeared on the agar plates. Each colony represents a unique genotype derived from an individual cell. *C. reinhardtii* is an isogamous species with two mating types (mt): mt− or mt+. A library of genotypes was established by random selection of 20 colonies of each mating type, collected from the agar plates by sterile loops. The mating type was determined by crossing the culture derived from each genotype with tester isolates.

The experimental populations were constructed by random selection of 10 different genotypes from the library. Each experimental population comprises a unique combination of 10 genotypes. The sexual populations were established by combining 5 mt− and 5 mt+ isolates, while the asexual populations were constructed by combining either 10 mt+ or 10 mt− isolates (with an equal proportion of populations comprising either mating type).

### 2.2 Selection experiment

#### Treatments

The experimental populations were subjected to a deteriorating environment, which comprised an increase of NaCl (hereafter referred to as salt) level in the medium in regular intervals, at three different rates. The salt was chosen as a stressor since previous experiments (Lachapelle and Bell, 2012; Lachapelle et al., 2015) demonstrated the detrimental effect of elevated salt concentration on *C. reinhardtii* populations, manifested through osmotic and ionic disbalance in cells (Lachapelle et al., 2015).

We manipulated the life cycle of *C. reinhardtii* by shifting between a stage during which all the populations were reproducing mitotically (hereafter referred to as the asexual stage) and a stage of sexual reproduction. The asexual stage consisted of 4 growth cycles (3-4 days each), during which the populations underwent approximately 4-6 rounds of mitotic divisions per growth cycle (9-12 doublings of cells). During this stage, salt level remained constant. After completion of the asexual stage, a sexual cycle has been initiated in all the populations, but completed only in the sexual populations, due to presence of both mating types (see ‘Sexual cycle’ section below for details). After completion of each sexual cycle, the asexual stage has been reinitiated, during which the experimental populations experienced an increased level of salt.

The three treatment groups were established depending on the rate of salt increase which the populations experienced: the populations in the gradual rate treatment group experienced a relatively slow increase of salt concentration (1 g/l per each asexual stage); the populations in the high rate treatment group experienced a relatively abrupt increase of salt concentration (3 g/l per each asexual stage); the moderate rate treatment group was subjected to an intermediate level of salt increase (2 g/l per each asexual stage). Each treatment group consisted of 72 populations.

The three treatment groups were established with respect to mode of reproduction: the asexual, obligate sexual and facultative sexual. The populations within both the obligate sexual and facultative sexual treatment groups were allowed to complete a number of rounds of sexual reproduction during the experiment. In the populations of the obligate sexual group, the gametes which failed to mate during the sexual cycle were eliminated by freezing. Thus, only sexually derived progeny were allowed the subsequent asexual stage. In the facultative sexual group, this step was omitted, resulting in mixed progeny derived from both the zygotes and unmated gametes. The asexual populations were reproducing entirely mitotically throughout the course of the experiment. Within each treatment group (72 populations each), 24 populations were subjected to one of the three rates of environmental deterioration. However, one of the genotypes sampled from the library was incorrectly assigned as a plus mating type, which was noted after the experiment had already commenced. Since the presence of two mating types may have resulted in zygote production in 14 asexual populations containing that genotype, these have been withdrawn from the subsequent analysis of the experiment, which included seven gradual rate populations, one moderate rate population and six high rate populations.

#### Cultivation and Transfer Procedure

All experimental populations were cultivated in 24-well microtiter plates in Bold’s basal medium (2 ml per culture), widely used for algal cultivation (Harris, 2009), under standard conditions (26°C, 100 μE illumination, constantly shaken at 180 rpm (3-mm rotation diameter), covered with sterile breathable membranes to prevent uneven evaporation across the plates and cross-contamination). A serial passage was performed after completion of each growth cycle by transferring 5 % of each population to a fresh medium.

#### Sexual Cycle

After completion of the asexual stage, sexual populations were allowed to undergo a sexual cycle, using the following protocol. All the populations were centrifuged at 5000 rpm for 10 minutes and re-suspended in nitrogen-free medium to initiate gametogenesis. The populations were then incubated in standard conditions (without shaking) under bright light for 24 hours to allow for mating of gametes and formation of zygotes. After given period, the microtiter plates containing the experimental populations were wrapped in aluminium foil and incubated in the darkness for additional 4-5 days to allow the zygotes to mature. Since sub-lethal stress may affect the mutation rate, the effects of sex may be confounded with the effects of mutations (Goho and Bell, 2000; Colegrave et al., 2002). Hence, the nitrogen starvation treatment was also applied to the asexual populations, but since they were made up of single mating types, no mating took place. After incubation in the darkness, the microtiter plates with mature zygotes were placed in a freezer for 4 hours at −20°C in order to eliminate unmated gametes. The zygotes develop the additional layers of cell wall during maturation in the darkness, thereby acquiring resistance to stressful conditions (Harris, 2009). The pilot experiments carried out in our laboratory revealed that this characteristic enables the zygotes to withstand low temperature. Since unmated gametes lack resistance to freezing, this procedure was omitted for the asexual and facultative sexual groups, and the transfer of both the produced zygotes and unmated gametes to the next asexual stage was allowed in the latter group. After the freezing procedure, the zygotes produced by the obligate sexual populations were transferred to the agar plates by sterile loops and incubated in bright light for two days to allow for germination and several rounds of mitotic divisions. The zygotes produced by the facultative sexual populations (if any) were transferred using the same method as for the obligate sexual populations, along with additional aliquot (of about 50-100 μl of culture) to prevent the population bottleneck if a low number of zygotes had been produced. The asexual populations were transferred to the agar plates by pipetting (an aliquot of 150 μl) and incubated for the same period as the sexual populations. After given interval of time, all the cultures were flooded with 4-5 ml of Bold’s basal medium (containing a higher salt concentration than during the previous asexual stage, depending on the treatment group) for approximately an hour. Population size of each experimental population propagated on agar was estimated by spectrophotometer (OD_750_) and the populations were diluted to the same optical density as before induction of sexual cycle. This experimental procedure ensured that the effects of sex were not confounded with a variation in population size. All experimental populations were then returned to the liquid medium by pipetting.

The mating of *C. reinhardtii* requires sufficiently dense populations to ensure the contact of gametes and thus, production of zygotes. Since different rates of environmental deterioration caused variation in the rate of mean population size decline (see the ‘Results„section) and thereby variation in population densities prior to induction of sexual cycles, an unequal number of sexual cycles were induced for each treatment group. Consequently, the sexual populations within the gradual rate, moderate rate and high rate treatment groups underwent ten, four and three rounds of sexual reproduction, respectively.

### 2.3 Data collection

We investigated the interplay of mode of reproduction and the rate of environmental deterioration by measuring the three parameters: a decline of mean population size before first extinctions occurred, the rate of extinction events and the probability of evolutionary rescue.

#### Population size dynamics

After every second growth cycle, each population was sampled (150 μl of the culture) and population size estimated by measuring optical density (OD_750_) using spectrophotometer. The measurements were carried out up to and including the 20^th^ growth cycle of each population, since optical density of sparse cultures could not have been estimated in a reliable way using this method.

#### Extinction dynamics

Each experimental population was visually inspected under inverted microscope (with 20 × magnification) before each transfer to potentially record an extinction event (absence of living cells). If no surviving cells were detected after observation under the microscope, a sample of the population (200 μl, approximately 10% of the culture) was transferred to the agar plate and incubated for 3-4 days in order to confirm an extinction event. The selection experiment continued until all experimental populations went extinct.

#### Evolutionary rescue

At the point which salt concentration had reached and surpassed the level which could potentially completely stop the growth of the ancestral populations (8 g/l, as demonstrated by Reynoso and de Gamboa, 1982; Moser and Bell, 2011), thereby increasing the risk of extinction, we initiated a parallel experiment, by sub-sampling experimental populations and continuing to propagate each in a selective medium comprising the same salt concentration as at the time of sub-sampling, by means of serial passaging. During this experiment, the salt concentration remained constant. The objective of this experiment was to evaluate whether evolutionary rescue had occurred, manifested through positive growth in stressful conditions. Simultaneously, the main selection experiment was continued by subjecting the main populations to increasing salt concentrations.

The sub-sampled populations (5% of the culture) were transferred to 24-well plates with the corresponding selective medium and propagated for 3-4 additional growth cycles. Population size was estimated spectrophotometrically (OD_750_) after each growth cycle. After the second growth cycle, a sample (20 μl) of each population was transferred to the corresponding selective medium (180 μl) in 96-well plate to estimate the growth rate of the populations. The optical density (OD_750_) of the populations propagated in 96-well plates was measured twice a day (after every 12 h), until the cultures reached a mid-log/stationary phase (after approximately 3.5 days). The growth rate of each population, estimated by calculating yield as a function of time, was a measure of fitness in stressful salt concentrations.

The growth parameters (population size and growth rate) of these populations were compared with those of the ancestral isolates, assayed in the medium with the corresponding salt concentration prior to commencing of the selection experiment (data not shown). The populations were scored as “rescued” if they showed:

1) clear positive growth between transfers in 24-well plates;

2) positive growth rate when assayed in 96-well plates;

3) significantly faster growth rate than that of the ancestors (compared for the same salt concentration).

## 3. Data analysis

The population size dynamics of each treatment group was analysed by fitting Two-way ANOVA with two factors as categorical independent variables: ‘mode of reproduction’ and ‘the rate of environmental deterioration’, both treated as fixed factors. The dependent continuous variable was the slope of the regression line representing mean decline of population size as a function of time (in growth cycles). The interaction between the factors was incorporated into the model.

The Kaplan-Meier survival curves of each treatment group were analysed by fitting Weibull regression model with two factors: ‘mode of reproduction’ and ‘the rate of environmental deterioration„. The extinction dynamics was analysed with respect to time (the number of growth cycles each population underwent prior to extinction) and salt concentration. The interaction between the factors was incorporated into the model.

The probability of evolutionary rescue within each treatment group was estimated by fitting a Generalised Linear Mixed Model (GLMM) with two factors as categorical independent variables: ‘mode of reproduction’ and ‘the rate of environmental deterioration, both treated as fixed factors. The continuous independent variable ‘salt concentration’ was treated as random factor. The binary response variable was ‘rescue (positive growth) / death (negative growth)’. The interaction between fixed factors was incorporated into the model.

Mean fitness of rescued populations (growth rate) was analysed by fitting a Linear Mixed Model (LMM) with two factors as categorical independent variables: ‘mode of reproduction’ and ‘the rate of environmental deterioration, both treated as fixed factors. The continuous independent variable ‘salt concentration’ was treated as random factor. The continuous response variable was the slope of the regression line representing mean increase of population size as a function of time. The interaction between fixed factors was incorporated into the model. The data was transformed using Box-Cox transformation method prior to the analysis.

All the analyses were carried out using R (R Core Team, 2017).

## 4. Results

### Population size dynamics

To estimate the effect of mode of reproduction and the rate of environmental deterioration on population size dynamics, we regularly monitored population size of experimental populations by measuring the optical density of the cultures after every second growth cycle. The measurements were carried out up to and including the 20^th^ growth cycle. We estimated that experimental populations of *C. reinhardtii* underwent approximately 100 generations of cell divisions during this interval of time.

Mode of reproduction significantly affected the rate of decline of mean population size (two-way ANOVA; F_2,201_ = 102.26, P < 0.00001). The highest decline of mean population size was recorded in the asexual group, having been by 42 % and 36 % higher than that of the obligate sexual and facultative sexual group, respectively (Figure 1A). Post-hoc Tukey’s HSD analysis revealed statistically significant difference (P < 0.00001) between the asexual group and both the obligate sexual and facultative sexual group. No significant difference between the sexual groups of populations was detected (P = 0.07).

**Figure 1.**
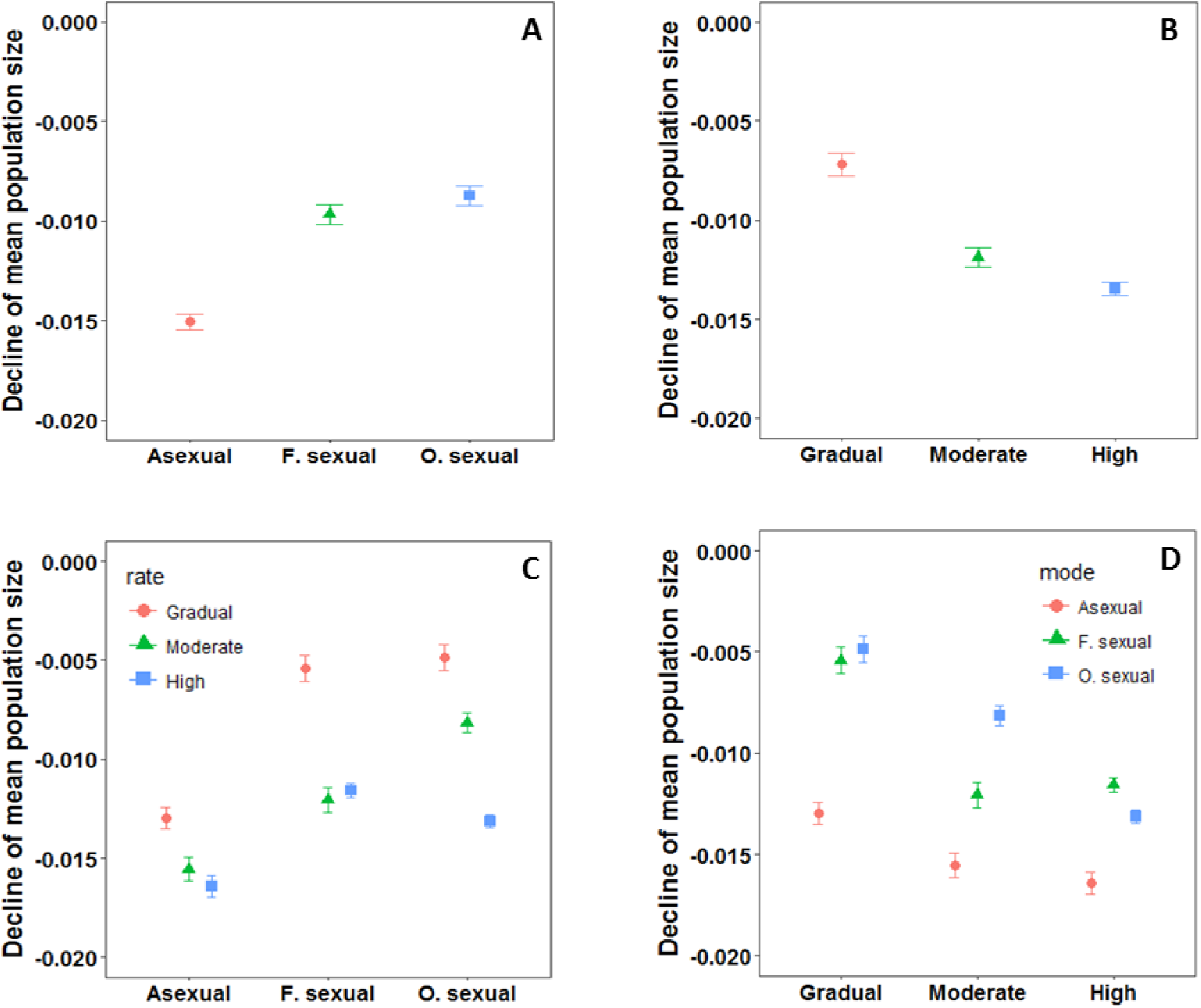
**The decline of mean population size: per mode of reproduction (A); per rate of environmental deterioration (B); per mode of reproduction with respect to the rate of environmental deterioration (C); per rate of environmental deterioration with respect to mode of reproduction**; the bars represent standard error of the mean.

The rate of environmental deterioration was positively correlated with the rate of decline of mean population size (two-way ANOVA; F_2,201_ = 102.99, P < 0.00001) (Figure 1B). The decline of population size was lowest in the treatment group subjected to the gradual rate of salt increase, and significantly lower than that of the treatment groups subjected to the moderate rate and high rate of environmental deterioration (by 39 % and 47 %, respectively) (Tukey’s HSD, P < 0.00001). The treatment group subjected to the moderate rate of salt increase had significantly lower decline of population size than that of the treatment group subjected to the high rate of salt increase (by 12 %) (Tukey’s HSD, P = 0.001).

There was a significant interaction between mode of reproduction and the rate of environmental deterioration (Two-way ANOVA; F_4,201_ = 10.15, P < 0.00001). While both sexual groups had a lower decline of population size relative to the asexual group with respect to all three rates of environmental deterioration (Tukey’s HSD, P < 0.005 for all pairwise comparisons) (Figure 1, C and D), the only significant difference between the sexual groups was detected under the moderate rate of environmental deterioration (Tukey’s HSD, P < 0.0001). The decline of mean population size of the moderate rate - obligate sexual group was lower by 32% than that of the moderate rate - facultative sexual group (Figure 2D).

**Figure 2.**
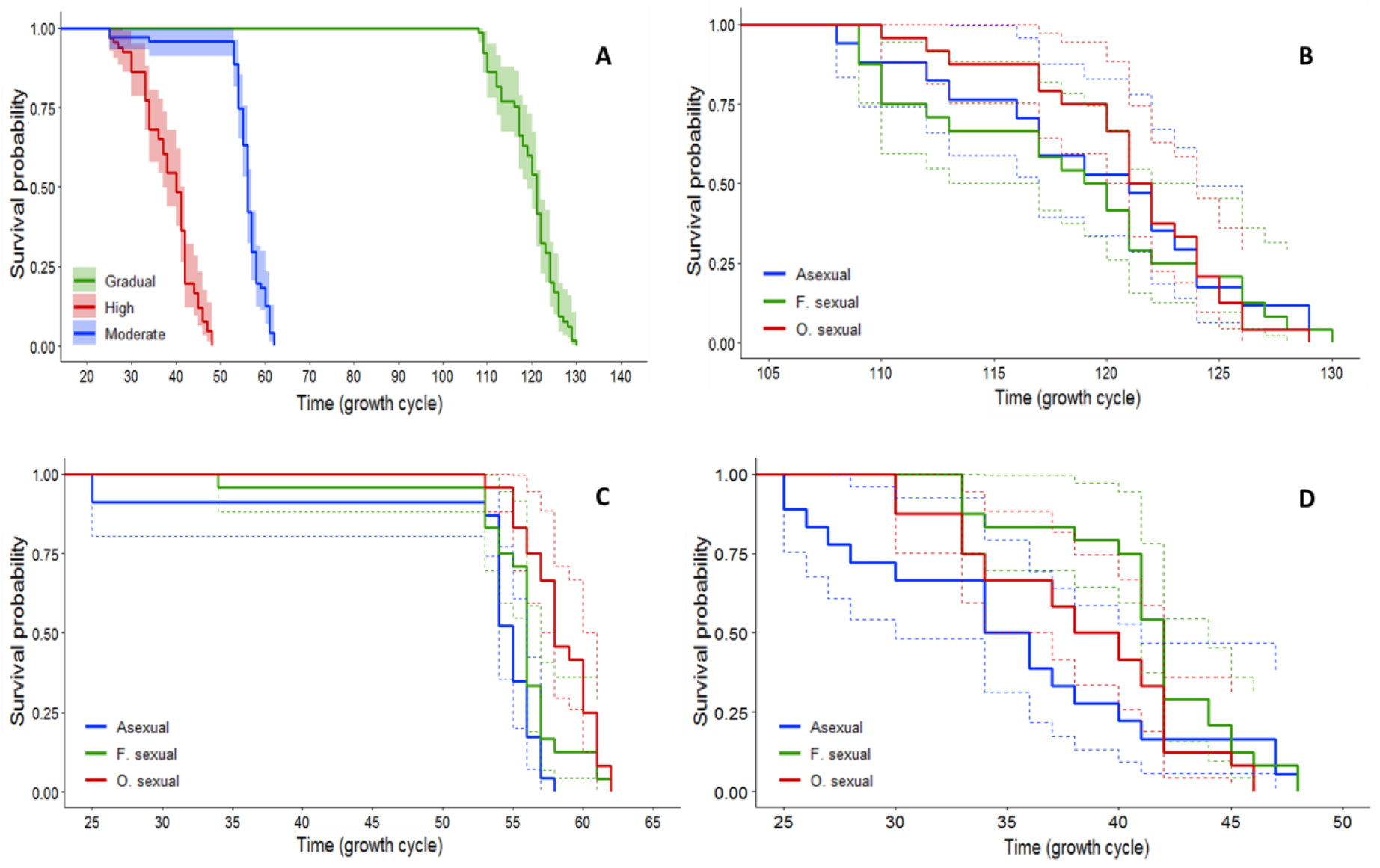
**The extinction dynamics of experimental populations: per rate of environmental deterioration (A); per mode of reproduction under the gradual rate of environmental deterioration (B); per mode of reproduction under the moderate rate of environmental deterioration (C); per mode of reproduction under the high rate of environmental deterioration (D)**; extinctions are plotted against time (the number of growth cycles that each population underwent prior to extinction); 95% confidence interval for each Kaplan-Meier survival curve is represented with the shaded area around the curves (A) or dotted lines (upper and lower boundary; B,C and D).

### Extinction dynamics

To investigate the effect of mode of reproduction and the rate of environmental deterioration on extinction dynamics, we monitored experimental populations after each growth cycle to record the potential extinction events. An experimental population was recorded as extinct if no cells were detected after visual inspection under the microscope and incubation of a culture sample on an agar plate. We fitted two Weibull regression models, to analyse the extinction dynamics against time (in growth cycles) and salt concentration, respectively. The best fit was obtained using extreme hazard function (for the model of the extinction dynamics with respect to time) and logistic hazard function (for the model of the extinction dynamics with respect to salt concentration), respectively.

The rate of extinctions of experimental populations against time in growth cycles was positively correlated with the rate of environmental deterioration (Weibull regression; χ^2^ = 805.81; df = 2; P < 0.00001) (Figure 2A). The extinction rate of the gradual rate group was significantly slower than that of both the high rate group (z = 56.23, P < 0.00001) and moderate rate group (z = 49.15, P < 0.00001). The extinction rate of the moderate rate group was significantly slower than that of the high rate group (z = 10.30, P < 0.00001). First extinctions were recorded within the moderate and high rate groups (during the 25^th^ growth cycle). Last experimental populations within the high rate group went extinct by the 48^th^ growth cycle (after approximately 225 generations) while the last extinctions within the moderate rate group occurred during the 62^nd^ growth cycle (after approximately 300 generations). First extinctions within the gradual rate group occurred during the 108^th^ growth cycle, while the last extinctions were recorded during the 130^th^ growth cycle (after at least 600 generations).

The main effect of mode of reproduction on extinction dynamics of experimental populations was not statistically significant (Weibull regression; χ^2^ = 3.51; df = 2; P = 0.17). However, the relative advantage of the individual levels within ‘mode of reproduction’ factor was contingent on the rate of salt increase, given the significant interaction between mode of reproduction and the rate of environmental deterioration (Weibull regression; χ^2^ = 10.58; df = 4; P = 0.03). While the Kaplan-Meier survival curves of each treatment group were not significantly different under the gradual and high rate of salt increase (Figures 2B and ^2^ = 10.58; df = 4; P = 0.03). While the Kaplan-Meier survival curves of each treatment group were not significantly different under the gradual and high rate of salt increase (2D, respectively), the obligate sexual group had significantly slower extinction rate relative to the asexual group (z = 2.3, P = 0.02) when subjected to the moderate rate of salt increase (Figure 2C).

The extinction dynamic of experimental populations was also analysed with respect to salt concentration upon each extinction event. The experimental populations had the highest probability of extinction under salt concentrations of 28 g/l (22% of all extinctions) and 30 g/l (29% of all extinctions) (Figure 3A). The rate of environmental deterioration significantly affected the extinction dynamic of experimental populations with respect to salt concentration (Weibull regression; χ^2^ = 18.74; df = 2; P = 0.00008) (Figure 3B). The extinction rate of the gradual rate group was significantly slower than that of the high rate group (z = 2.79; P = 0.005) and marginally slower than that of the moderate rate group (z = 1.85; P = 0.06); no significant difference was detected between the extinction dynamics of the moderate rate and high rate groups (z = 1.26; P = 0.2). First extinctions within the gradual rate group occurred under salt concentration of 27 g/l, 63% of the populations reached and 31% survived 30 g/l, while the last populations went extinct while being subjected to 32 g/l of salt concentration. First populations within the moderate rate group went extinct under 12 g/l of salt concentration, followed by a long interval (between 12 g/l and 26 g/l) during which only a single extinction occurred. The majority of populations (96%) went extinct within a relatively brief interval of time which corresponded to salt concentrations between 26 g/l and 30 g/l; 20% of populations reached, but none survived 30 g/l of salt concentration. First extinctions within the high rate group occurred under salt concentration of 18 g/l, 32% of the populations went extinct by the time which corresponded to salt concentration of 24 g/l, 45% of the populations reached and 20% survived 30 g/l. The last extinctions in this treatment group occurred under salt concentration of 36 g/l (see Figure 3D for extinction dynamics of all treatment groups, with respect to salt concentration).

**Figure 3.**
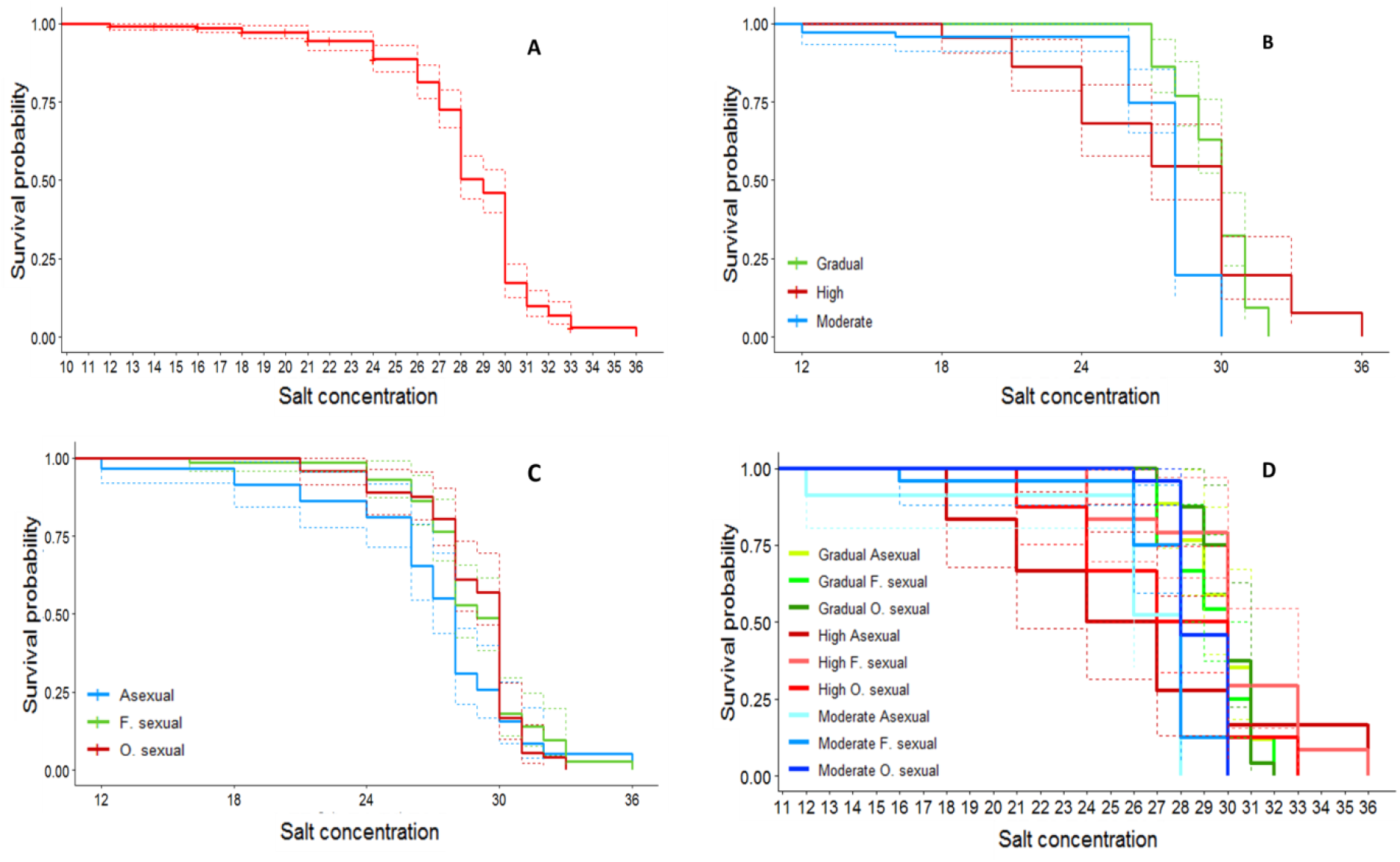
**The extinction dynamics of experimental populations: total probability of survival of each salt concentration (A); per rate of environmental deterioration (B); per mode of reproduction (C); with respect to both mode of reproduction and the rate of environmental deterioration (D)**; extinctions are plotted against the salt concentration upon each extinction event; 95% confidence interval for each Kaplan-Meier survival curve is represented with the dotted lines (upper and lower boundary).

Mode of reproduction significantly affected the extinction rate with respect to salt concentration (Weibull regression; χ^2^ = 10.87; df = 2; P = 0.004). The extinction rate of the populations within the obligate sexual group was significantly slower than that of the asexual group (z = 3.01; P = 0.002) and marginally slower relative to that of the facultative sexual group (z = 1.78; P = 0.07) (Figure 3C). No difference was detected between the extinction dynamics of the asexual and facultative sexual groups (z = 1.25; P = 0.21). There was a significant interaction between mode of reproduction and the rate of environmental deterioration on the extinction rate with respect to salt concentration (Weibull regression; χ^2^ = 22.94; df = 4; P =0.0001). The beneficial effect of the facultative sexual group significantly increased under the high rate of environmental deterioration, as reflected in slower extinction rate relative to both the asexual and obligate sexual groups (Figure 3D). The magnitude of the effect of facultative sexual mode of reproduction (on extinction rate) was significantly different than under both the gradual rate (z = 2.86; P = 0.004) and moderate rate (z =3.34; P = 0.0008) of salt increase.

### Evolutionary rescue

In parallel with propagating the main experimental populations in the environment with increasing concentration of salt, a second parallel study was carried out. Before being subjected to the next step of salt increase, each experimental population was sub-sampled and propagated for additional 3-4 growth cycles in the same level of salt as at the time of sub-sampling, while the growth rate (mean fitness) of these populations was tested against that of the ancestral populations. The objective of this study was to investigate whether evolutionary rescue had occurred during the main selection experiment.

We recorded 137 evolutionary rescue events in total, out of 475 assayed populations (29%). Mode of reproduction significantly affected the probability of evolutionary rescue (GLMM; χ^2^ = 8.36; df = 2; P = 0.015). The populations within the obligate sexual group had a higher probability of evolutionary rescue relative to both the asexual and facultative sexual groups (z = 2.80 and z = 2.67, respectively; P < 0.008, for both pairwise comparisons) (Figure 4A). No significant difference between the asexual and facultative sexual groups was detected.

**Figure 4.**
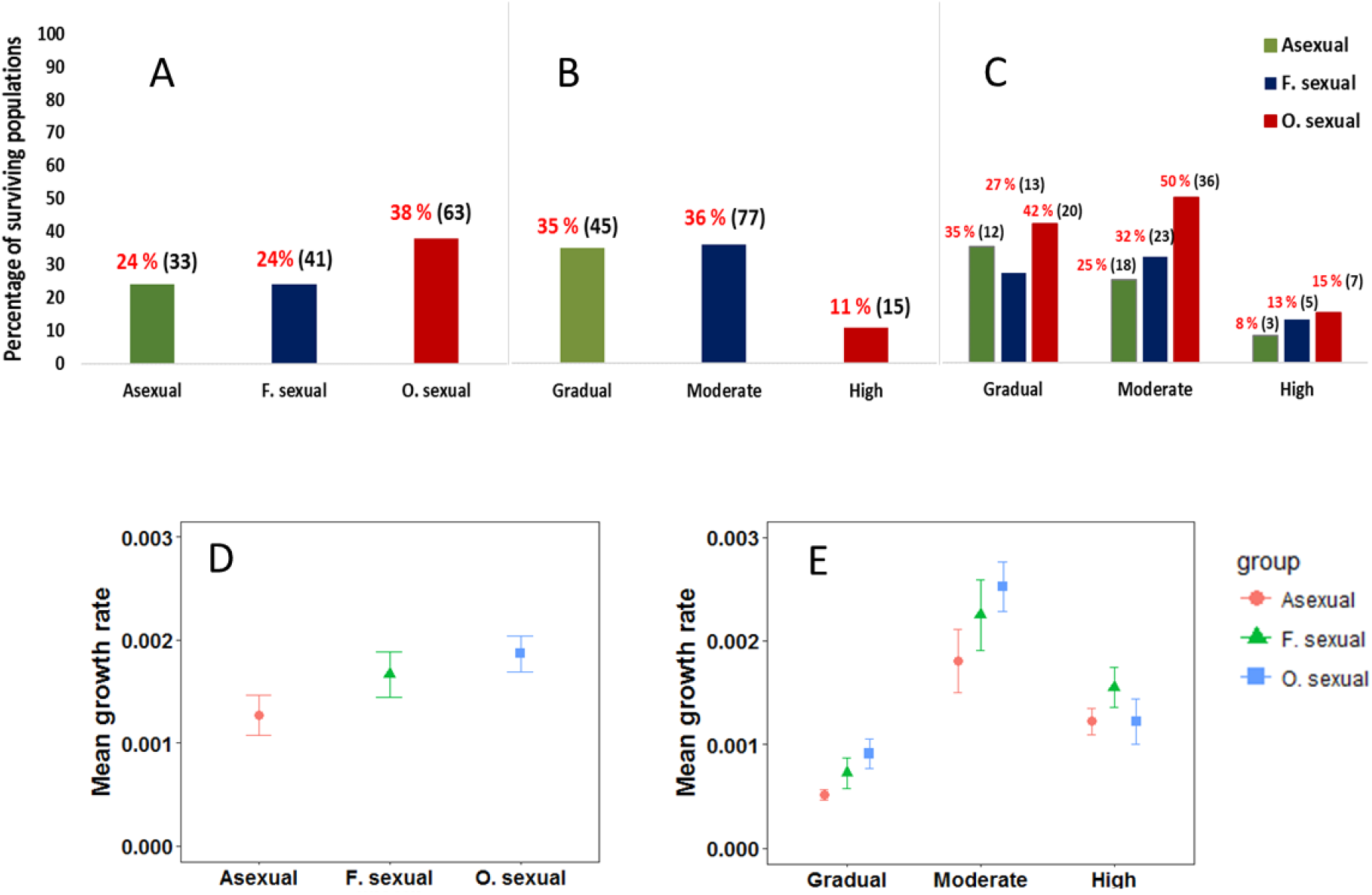
**The probability of evolutionary rescue of experimental populations: per mode of reproduction (A); per rate of environmental deterioration (B); per rate of environmental deterioration with respect to mode of reproduction (C)**; percentage of rescued experimental populations per each treatment group is presented above each bar plot, with the number of rescued populations in brackets; **mean fitness of rescued populations (growth rate): per mode of reproduction (D); per rate of environmental deterioration with respect to mode of reproduction (E)**; evolutionary rescue of the populations within the graduate rate group was tested for salt concentrations of 16 g/l and 17 g/l, respectively; evolutionary rescue of the populations within the moderate rate group occurred during exposure to salt concentrations of 8 g/l, 10 g/l and 12 g/l, respectively; evolutionary rescue of the populations within the high rate group was detected during exposure to salt concentrations of 9 g/l and 12 g/l, respectively; the bars represent standard error of the mean.

The probability of evolutionary rescue was significantly affected by the rate of environmental deterioration (GLMM; χ^2^ = 22.99; df = 2; P < 0.00001). The populations within the gradual rate and moderate rate treatment groups had significantly higher probability of evolutionary rescue relative to the high rate treatment group (z = 3.91 and z = 4.31, respectively; P < 0.0001, for both pairwise comparisons) (Figure 4B). No significant difference between the gradual rate and moderate rate groups was detected. Within the gradual rate group, 17 g/l was the maximal salt concentration for which evolutionary rescue was recorded. By contrast, the maximal salt concentration survived by any population within both the moderate and high rate groups was 12 g/l.

No significant interaction was detected between mode of reproduction and the rate of environmental deterioration (GLMM; χ^2^ = 1.08; df = 4; P = 0.9). The populations within the obligate sexual group had the highest probability of evolutionary rescue irrespective of the rate of salt increase (Figure 4C).

Mean fitness of rescued populations was significantly affected by mode of reproduction (LMM; F_2,134_ = 6.29; P = 0.002). Mean growth rate of the obligate sexual populations was significantly faster than that of the asexual populations (t = 3.43; df =94; P = 0.0009) and marginally faster than that of the facultative sexual populations (t = 1.98; df =102; P = 0.05) (Figure 4D). There was no significant difference between mean growth rate of the asexual and facultative sexual populations. No significant interaction between mode of reproduction and the rate of environmental deterioration on fitness of rescued populations was detected (LMM; F_4,134_ = 0.62; P = 0.65) (Figure 4E).

## 5. Discussion

The objective of this study was to investigate the interplay of mode of reproduction and the rate of environmental deterioration on extinction dynamics and the probability of evolutionary rescue of *C. reinhardtii* experimental populations, in environment deteriorating at different rates. The main prediction of the experiment was an increase of relative advantage of sexual populations under higher rates of environmental deterioration.

Overall, sexual mode of reproduction was beneficial. Sex significantly reduced both the rate of population size decline and the extinction rate with respect to salt concentration, and increased the probability of evolutionary rescue. The sexual groups combined had 39% lower decline of population size relative to the asexual group. The obligate sexual group had significantly slower extinction rate than that of the asexual group, as reflected in higher number of surviving populations during exposure to salt concentrations between 12 g/l (under which first extinctions had occurred) and 30 g/l. Finally, the obligate sexual group had the highest proportion of evolutionary rescue events and the fastest growth rate of the rescued populations.

We detected the interaction between mode of reproduction and the rate of environmental deterioration on both the rate of population size decline and the extinction rate (with respect to both time in growth cycles and salt concentration). Under the moderate rate of environmental deterioration, the obligate sexual group had the lowest population size decline and significantly slower extinction rate (with respect to time in growth cycles) than that of the asexual group. The facultative sexual group had the slowest extinction rate (with respect to salt concentration) under the high rate of environmental deterioration.

This experiment demonstrated that the sign of the effect of sex remains constant irrespective of the rate of environmental deterioration (the advantage of at least one or both sexual groups relative to the asexual group), as predicted. Furthermore, we detected that the relative advantage of sex (slower extinction rate) increased with the rate of environmental deterioration. While no sexual group had significant advantage over the asexual group under the gradual rate of environmental deterioration, the sexual groups had significantly slower extinction rate under the moderate rate (obligate sexual) and high rate (facultative sexual) of environmental deterioration. One of the possible explanations for this result is that the adaptive walk of the asexual populations in a gradually changing environment was as efficient as of the sexual populations, hence the relative advantage of sex diminished. The relative advantage of sex with respect to population size decline and the probability of evolutionary rescue was relatively constant and did not depend on the rate of environmental deterioration.

The extinction dynamic of the sexual populations depended on the rate of environmental deterioration. While the obligate sexual group had marginal relative advantage over the facultative sexual group under the gradual and moderate rate of environmental deterioration, facultative sexuality was a more beneficial strategy under the high rate of environmental deterioration. The potential explanation for this shift of relative advantage between the sexual groups is a high demographic cost of adaptation imposed by the high rate of environmental deterioration. The obligate sexuality may have generated higher diversity of genotypes (genetic variance for fitness increased) relative to facultative sexuality during this treatment, but at a cost of mean fitness decline. This effect of sexual reproduction (recombination load) manifested during adaptation to changing environments has been previously demonstrated in experimental studies (Greig et al., 1998; Colegrave et al., 2002; Kaltz and Bell, 2002; Becks and Agrawal, 2012). The obligate sexual populations may have adapted with a simultaneous decline of population size, due to a strong selection which operated under the high rate of environmental deterioration, coupled with a genetic cost of sex (recombination load). By contrast, the facultative sexual populations may have been less well adapted, but suffered less pronounced population size decline. We recorded a lower population size decline in the facultative sexual group relative to the obligate sexual group under the high rate of environmental deterioration, though this effect was not statistically significant. In addition, the populations within the facultative sexual group comprised a fraction of individuals (derived from unmated gametes) which had not undergone sexual reproduction. The presence of these individuals may have compensated for recombination load, arose as a consequence of mating and zygote production of other fraction of the population, since mean fitness of this fraction remained relatively constant (thereby reducing recombination load of the population as a whole).

While the relative advantage of the obligate or facultative sexual group with respect to the extinction rate varied, depending on the rate of salt increase, obligate sexuality was advantageous with respect to the probability of evolutionary rescue, irrespective of the rate of environmental deterioration. The obligate sexual populations were more likely to be rescued than those in other treatment groups, and this effect was consistent under all three rates of environmental deterioration. Moreover, rescued obligate sexual populations also had the fastest growth rate. Both the highest occurence of rescue events and highest fitness of rescued populations within the obligate sexual group could be attributable to higher diversity of genotypes favourable in stressful environments generated by obligate sex. In addition, mean population size decline was lowest in this treatment group (though only marginally significant relative to the facultative sexual group). Since supply of beneficial muations is higher in larger populations (Samani and Bell, 2010), the obligate sexual populations could have been more likely to adapt through *de novo* beneficial mutations. Our study corroborated the results of Lachapelle and Bell (2012), who found that obligate sexuality (coupled with high genetic diversity) positively affects the probability of evolutionary rescue and adaptation to stressful environments (elevated salt concentrations), and demonstrated that this beneficial effect does not depend on the rate of environmental deterioration.

Taken together, the results of our experiment provide evidence that the probability of survival of adapting populations may depend on their mode of reproduction and that sex may prolong survival in stressful (and even lethal) conditions. However, this effect will be less pronounced in a gradually changing environment, because the asexual populations may keep up with the pace of environmental change as efficiently as the sexual populations. If environmental change proceeds at higher rates, different modes of sexuality may be beneficial under different rates of environmental change: obligate sexuality may be advantageous under the intermediate rates of change (and moderate strength of selection), while facultative sexuality will be favoured under higher rates of change (more intense selection). Furthermore, to the best of our knowledge, our study is the first to demonstrate that the beneficial effect of sex on adaptation to stressful environments does not depend on the rate of environmental change and that obligate sexuality maximizes the probability of evolutionary rescue.

Given the dependence of the effects of sex on the rate of environmental deterioration and widely the recognized issue of global change (for example, Falkowski et al., 1994; Chivian and Bernstein, 2008; Barnosky et al., 2012), which may proceed at a rate unprecedented in the Cenozoic (Barnosky et al., 2012), one of the logical question that our findings may raise is: will the probability of survival of the extant species be increased by sex? In nature, many organisms are characterized by obligate sexual reproduction (for example, birds and mammals). The obligate sexual organisms tend to have longer generation time and thus, lower reproductive output relative to the organisms that reproduce mainly asexually. If standing genetic variation of these species declines due to reduction of population size (a likely consequence of global change), we could predict a slower rate of adaptation and potentially a higher extinction risk for these organisms. The obligate sex could increase the response of selection through well-understood mechanisms, and thus prolong population persistence. However, given that primary response to a long-term environmental change involves *de novo* beneficial mutations, which are unlikely to arise fast enough in slower reproducers, we could predict the relatively modest long-term effects of obligate sex, and phenotypic plasticity (for example, migration or behavioural change) as the main adaptive response of the obligate sexual populations. Thus, survival of current (anthropogenically induced) environmental change may be more likely for small, fast reproducing, facultative sexual organisms, with larger population sizes and higher supply rate of beneficial mutations (e.g. microorganisms).

## Acknowledgements

NP has been funded by the Darwin Trust of Edinburgh.

## Data accessibility

Data will be available through the Dryad Digital Repository (datadryad.org).

